# Intravenous edaravone plus therapeutic hypothermia offers limited neuroprotection in the hypoxic-ischaemic newborn piglet

**DOI:** 10.1101/2020.02.25.964288

**Authors:** Satoshi (Hamano) Yamato, Shinji Nakamura, Yinmon Htun, Makoto Nakamura, Wataru Jinnai, Yasuhiro Nakao, Tsutomu Mitsuie, Kosuke Koyano, Takayuki Wakabayashi, Aya (Hashimoto) Morimoto, Masashiro Sugino, Takashi Iwase, Sonoko (Ijichi) Kondo, Saneyuki Yasuda, Masaki Ueno, Takanori Miki, Takashi Kusaka

## Abstract

Therapeutic hypothermia is a standard therapy for neonatal hypoxic-ischaemic encephalopathy. One potential additional therapy is the free radical scavenger edaravone (3-methyl-1-phenyl-2-pyrazolin-5-one). To compare the neuroprotective effects of edaravone plus therapeutic hypothermia with those of therapeutic hypothermia alone after a hypoxic-ischaemic insult in the newborn piglet, anaesthetized piglets were subjected to 40 min of hypoxia (3–5% inspired oxygen) and cerebral ischaemia was assessed by cerebral blood volume. Body temperature was maintained at 38.5 °C in the normothermia group (NT, n = 8) and at 34 °C (24 h after the insult) in the therapeutic hypothermia (TH, n = 7) and therapeutic hypothermia plus edaravone (3 mg intravenous every 12 h for 3 days after the insult; TH+EV, n = 6) groups under mechanical ventilation. Five days after the insult, the mean (standard deviation) neurological scores were 10.9 (5.7) in the NT group, 17.0 (0.4) in the TH group (*p* = 0.025 vs. NT) and 15.0 (3.9) in the TH+EV group. The histopathological score of the TH+EV group showed no significant improvement compared with that of the other groups. In conclusion, edaravone plus therapeutic hypothermia had no additive neuroprotective effects after hypoxia-ischaemia in neurological and histopathological assessments.

## Introduction

Neonatal hypoxic-ischaemic encephalopathy (HIE) is a clinically significant disorder with long-term morbidity [1,2]. Therapeutic hypothermia (TH) has been used to limit brain damage in term newborns, although it is unable to fully rescue neurological outcomes [3,4]. Therefore, other therapies that augment the neuroprotection of TH in neonatal HIE are needed.

Edaravone (3-methyl-1-phenyl-2-pyrazolin-5-one) is a synthetic free radical scavenger used as a neuroprotective drug. Edaravone has promising activities as an antioxidative radical scavenger, quenching hydroxyl radical (HO•) and inhibiting both HO•-dependent and •OH-independent lipid peroxidation [5,6]. The oxidative stress that occurs after an ischaemic stroke produces reactive oxygen species (ROS) such as hydrogen peroxide (H_2_O_2_), HO• and superoxide anion radical (O_2_^−^) that bring about membrane lipid peroxidation and vascular compartment endothelial cell injury [7]. Edaravone regulates numerous signalling pathways to delay neuronal death [8], reduce cerebral oedema, counteract microglia-induced neurotoxicity [9] and decrease long-term inflammation. Accordingly, many adult animal and human studies have reported the efficacy of edaravone in treating brain injury due to cerebral infarction [5–8,10]. Indeed, Radicut®, a brand of edaravone marketed by Mitsubishi Tanabe Pharma Corporation (Tokyo, Japan), has been approved in Japan to treat acute ischaemic stroke patients presenting within 24 h of the attack since 2001.

In newborns, oxidative stress also plays an important role in the biochemical cascade that leads to neuronal cell death after hypoxia-ischaemia (HI) [11,12]. Neonatal rat studies have revealed that edaravone reduces oxidative stress and improve outcomes after HI [13–16]. Ni et al. [17] had already reported the efficacy of edaravone for neuroprotection in a 3–7-day-old piglet model, but they did not use newborn piglets and those treated with TH. In this regard, assessment of the efficacy of edaravone in a perinatal large-animal model would be useful for preclinical evaluation. In this study, we used a previously developed perinatal asphyxia model of newborn piglets that survived 5 days after the HI insult [18].

Edaravone may become an additional therapy to TH for neonatal HI encephalopathy in humans. To our knowledge, no studies have revealed the efficacy of edaravone combined with TH in a large perinatal animal. We thus compared the neuroprotective effects of edaravone plus TH with those of TH alone after a HI insult in our newborn piglet model of perinatal asphyxia using intravenous edaravone administered for 3 days combined with TH for 24 h after resuscitation.

## Methods

### Animals

The study protocol was approved by the Animal Care and Use Committee for Kagawa University (15070-1) and in accordance with the Animal Research: Reporting In Vivo Experiments guidelines. Thirty-eight piglets obtained within 24 h of birth and weighing 1.63–2.10 kg were used in this study.

Twenty-one newborn Camborough piglets (Daiwa Chikusan, Kagawa, Japan) were initially anaesthetized with 1–2% isoflurane (Forane® inhalant liquid; Abbott Co., Tokyo, Japan) in air using a facemask. Each piglet was then intubated and mechanically ventilated with an infant ventilator. The umbilical vein and artery were cannulated with a 3- or 4-FG neonatal umbilical catheter (Atom Indwelling Feeding Tube for Infants; Atom Medical Co., Tokyo, Japan); the umbilical vein catheter was at a site 5 cm in depth from the incision and the umbilical artery catheter was at a site 15 cm in depth from the incision for blood pressure monitoring and blood sampling, respectively. After cannulation, the piglets were anaesthetized with fentanyl citrate at an initial dose of 10 µg/kg followed by infusion at 5 µg/kg/h and were paralysed with pancuronium bromide at an initial dose of 100 µg/kg followed by infusion at 100 µg/kg/h. Maintenance solution (electrolytes plus 2.7% glucose KN3B; Otsuka Pharmaceutical Co., Tokyo, Japan) was infused continuously at a rate of 4 mL/kg/h via the umbilical vein (glucose was infused at a rate of 2 mg/kg/min). Arterial blood samples were taken at critical points and when clinically indicated throughout the experiment. Each piglet was then placed in a copper mesh-shielded cage under a radiant warmer to maintain a rectal temperature of 39.0 ± 0.5 °C. Inspired gas was prepared by mixing O_2_ and N_2_ gases to obtain the oxygen concentrations required for the experiment. Ventilation was adjusted to maintain PaO_2_ and PaCO_2_ within their normal ranges. Arterial blood pressures were measured and recorded via the umbilical arterial catheter.

### Amplitude-integrated electroencephalography monitoring

The amplitude-integrated electroencephalography (aEEG) measuring device used was the Nicolet One (Cardinal Health, Inc., Dublin, OH), which displays the signal on a semi-logarithmic scale at low speed (6 cm/h). Measurements were recorded at 1-s intervals. Gold-plated electrode discs were placed at the P3 and P4 positions (corresponding to the left and right parietal areas on the head). Low-amplitude electroencephalography (LAEEG) was defined as a maximum amplitude < 5 μV.

### Hypoxic insult

Because the details were reported in our previous studies [19,20], only an outline of the HI insult protocol is presented here. Hypoxia was induced by reducing the inspired oxygen concentration of the ventilator to 4% after at least 120 min of stabilisation from the initial anaesthetic induction. If required to obtain a low-amplitude aEEG pattern (< 5 µV), the inspired oxygen concentration was further reduced to 2%. From the beginning of the LAEEG, the insult was continued for 30 min. FiO_2_ was decreased (1% decrements) or increased (1% increments) during the insult to maintain LAEEG, HR > 130 beats/min and MABP > 70% of baseline. Upon the reinstatement of LAEEG, HR or MABP, the FiO_2_ was returned to 4% during the first 20 min of the insult. For the final 10 min of the 30-min insult, if the MABP exceeded 70% of the baseline, hypotension was induced by decreasing the FiO_2_ until the MABP declined to below 70% of the baseline. During the HI insult, the cerebral blood volume (CBV) was continuously monitored using time-resolved spectroscopy (TRS-21; Hamamatsu Photonics K.K., Hamamatsu, Japan); we calculated the change in the CBV as described previously [18]. When the CBV fell to 30% of the height between the peak and baseline during the final 10 min of the insult, resuscitation was started. Otherwise, if the change in CBV was kept above it, the HI lasted for 10 min. Hypoxia was terminated by resuscitation with 100% oxygen. A base excess below –5.0 mEq/L was corrected as much as possible by sodium bicarbonate infusion to maintain a pH of 7.3–7.5. After 10 min of 100% FiO_2_, the ventilator rate and FiO_2_ were gradually reduced to maintain a SpO_2_ of 95–98%.

### Post-insult treatment

The piglets in all groups received mechanical ventilation for 24 h after resuscitation. Immediately after HI, the piglets in the TH+EV group (n = 6) were given intravenous edaravone (Radicut®; 3 mg/kg) every 12 h for 3 days after the insult. Because Yasuoka et al. [14] reported that repeated injection of edaravone (3 mg/kg) every 12 h for 7 days after HI insult decreased both apoptosis and necrosis in the neonatal rat, we chose the same dose in this study. The NT (n = 8) and TH (n = 7) groups were given an intravenous injection of saline for 3 days. Piglets in the NT group were maintained after resuscitation at 39 ± 0.5 °C. under a radiant heater. In the TH and TH+EV groups, whole-body hypothermia was achieved using a cooling blanket (Medicool; MAC8 Inc., Tokyo, Japan) after resuscitation. The piglets were cooled to 33.5 ± 0.5 °C for 24 h and then rewarmed at 1 °C/h using a blanket. The oesophageal temperature was used as the measure of body temperature. The temperature of the incubator was maintained at 28–32 °C. Once the piglets were weaned off the anaesthesia and ventilator and extubated, they were allowed to recover and were maintained for 5 days in the incubator. Piglets were fed 50–80 mL artificial animal milk via a nasogastric tube every 6 h. The presence of seizures was recognised clinically as rhythmic pathologic movements (cycling) and tonic postures sustained between cycling episodes. If seizures occurred, the piglet was treated with phenobarbital (20 mg/kg) via intramuscular injection. If seizures persisted, the piglet was treated with two successive anticonvulsant doses. If seizures persisted after two successive anticonvulsant doses, the piglet was euthanized.

### Neurological score

As reported by Thoresen et al. [21], we determined a nine-item (three-step) neurology score (0 abnormal, 1 mildly abnormal, 2 normal; total score range, 0–18) before intubation and 24 h, 48 h, 72 h and 5 days after the insult. The nine items examined during the neurological assessment were respiration, consciousness, orientation, ability to walk, forelimb control, hind limb control, limb tone, overall activity, sucking, vocalisation and presence of pathological movements. We also examined when piglets started to walk after the insult.

### Histology

After the 5-day period, the animals were initially anaesthetized with isoflurane, and the brain of each animal was perfused with 0.9% saline and 4% phosphate-buffered paraformaldehyde. Brain tissue was histologically evaluated and irregularities were graded according to a histopathology grading scale for a piglet model of post-hypoxic encephalopathy [21]. Coronal blocks of cortical grey matter, white matter, the hippocampus and the cerebellum were embedded in paraffin and cut with a microtome at 4 µm. The sections were stained with hematoxylin and eosin. At regular intervals of 100 µm, five sections each of samples were examined by investigators (M.U., S.N., S.Y. and Y.H.) who were blinded to all clinical information. The extent of damage in each of the five regions was graded in 0.5-unit intervals on a nine-step scale that ranged from 0.0 to 4.0.

### Data analysis

GraphPad Prism 5J (GraphPad Software, La Jolla, CA) was used for all statistical analyses. Blood sample and physiological data were compared among the three groups using two-way analysis of variance (ANOVA), whereas neurological score and histopathological score were compared among the three groups using one-way ANOVA. A *p* value < 0.05 was considered significant.

## Results

### Physiological results

In our animal model, heart rate (HR) and mean arterial blood pressure (MABP) increased during the approximate 40-min period of the hypoxic insult before subsequently decreasing, with similar values found in all groups (Fig 1).

**Fig 1.**
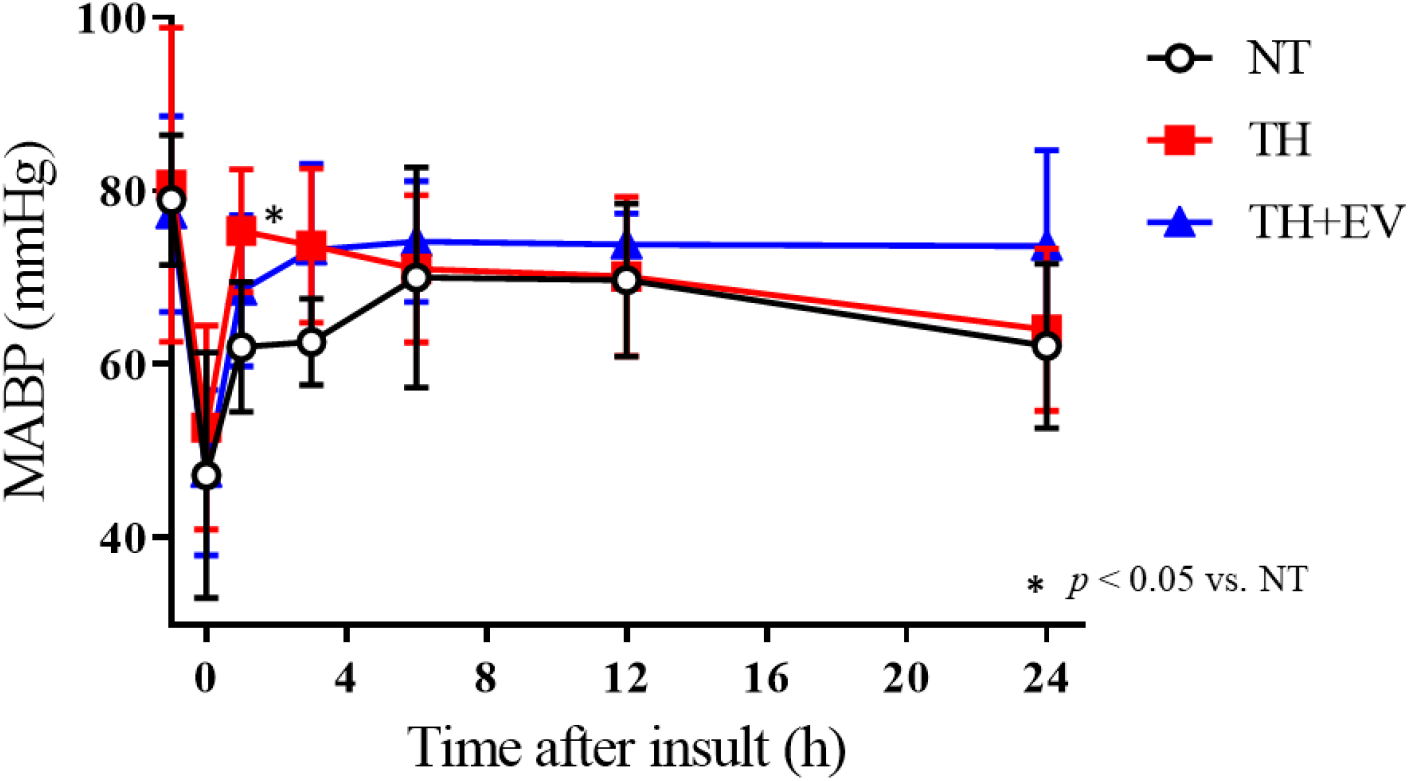
Time course of heart rate (HR) in three groups (NT, n=8; TH, n=7, TH+EV, n=6) at baseline, end of insult (0h), 3, 6, 12, 24 h after insult.

MABP was higher within 3 h after the insult in the TH (1 h, *p* < 0.05 vs NT) and TH+EV groups compared with the NT group, whereas HR was significantly lower in the TH (1, 3, 6, 12 h) and TH+EV (3, 6 h) groups than in the NT group after the insult (Fig 2). After resuscitation, arterial blood pH (pHa), partial pressure of arterial blood O_2_ (PaO_2_) and partial pressure of arterial blood CO_2_ (PaCO_2_) recovered to baseline values within 1 h in each group (Table 1). Some significant differences in base excess were detected among the groups at 6 and 12 h after resuscitation, but these differences were not considered physiologically significant. Hypothermia produced a transient but highly variable hyperglycaemic response at 6, 12 and 24 h after the insult. In addition, the TH-EV group showed more hyperglycaemia than the HI-TH group.

**Table 1.** Mean (SD) values of arterial pHa, PaCO_2_, PaO_2_, BE, blood glucose and lactate before, at the end of (0 h) and 1, 3, 6, 12 and 24 h after hypoxic-ischaemic insult in the three groups

| Parameters | Group | Baseline | 0 | 1 | 3 | 6 | 12 | 24 |
| --- | --- | --- | --- | --- | --- | --- | --- | --- |
| pHa | NT | 7.42 (0.05) | 6.85 (0.10) | 7.30 (0.06) | 7.48 (0.03) | 7.46 (0.05) | 7.46 (0.05) | 7.50 (0.06) |
|  | TH | 7.42 (0.11) | 6.79 (0.11) | 7.22 (0.12) | 7.42 (0.10) | 7.44 (0.05) | 7.44 (0.06) | 7.42 (0.03) |
|  | TH+EV | 7.45 (0.07) | 6.85 (0.12) | 7.20 (0.06)* | 7.40 (0.08) | 7.39 (0.04) | 7.45 (0.08) | 7.44 (0.06) |
| PaCO<sub>2</sub> (mmHg) | NT | 46.9 (4.9) | 34.5 (11.3) | 42.2 (6.6) | 42.5 (4.14) | 47.5 (6.4) | 45.1 (5.0) | 39.3 (3.3) |
|  | TH | 43.3 (12.5) | 42.5 (12.9)<sup>##</sup> | 42.2 (8.7) | 41.5 (8.0) | 42.2 (8.0) | 41.1 (7.7) | 37.4 (5.5) |
|  | TH+EV | 40.1 (6.4) | 29.5 (8.6) | 44.2 (5.3) | 41.0 (8.0) | 42.2 (7.4) | 40.0 (5.8) | 37.6 (6.0) |
| PaO<sub>2</sub> (mmHg) | NT | 89.6 (11.5) | 21.0 (11.9) | 94.5 (28.4) | 88.0 (18.1) | 89.2 (10.0) | 85.8 (15.2) | 89.7 (15.1) |
|  | TH | 99.4 (15.4) | 19.3 (7.2) | 110.3 (20.5) | 81.4 (21.2) | 81.7 (25.3) | 81.2 (25.7) | 78.6 (22.0) |
|  | TH+EV | 96.8 (22.8) | 20.2 (5.8) | 108.1 (22.8) | 86.0 (14.8) | 81.5 (13.4) | 99.4 (20.7) | 110.5 (34.4)** |
| BE (mmol/mL) | NT | 5.8 (2.4) | -26.0 (5.0) | -5.5 (3.9) | 7.8 (1.6) | 8.6 (1.6) | 7.2 (3.4) | 6.8 (2.7) |
|  | TH | 2.9 (3.5) | -27.6 (3.5) | -10.0 (4.9)* | 1.8 (4.2)** | 4.2 (3.2)* | 3.2 (3.4) | 0.7 (3.9)*** |
|  | TH+EV | 4.2 (3.1) | -26.0(2.4) | -10.1 (2.9)* | 2.6 (3.1)* | 4.6 (2.5)* | 3.5 (3.3) | 1.0 (4.2)* |
| Blood glucose<br>(mg/dL) | NT | 146.0 (19.4) | 253.5 (73.2) | 237.0 (57.0) | 218.1 (65.7) | 183.9 (56.8) | 195.5 (72.6) | 115.6 (40.1) |
|  | TH | 152.4 (42.3) | 226.6 (112.1) | 259.4 (75.1) | 261.0(64.9) | 229.0 (71.4) | 236.4 (66.2) | 196.3 (53.4) |
|  | TH+EV | 159.3 (34.7) | 296.8 (82.0) | 294.5 (76.9) | 311.8 (85.7)* | 298.0 (55.4)** | 293.3 (85.9)* | 186.2 (58.3) |
| Lactate (mg/dL) | NT | 18.0 (4.4) | 227.1 (28.0) | 122.5 (30.3) | 37.5 (11.6) | 27.8 (9.9) | 36.9 (11.8) | 34.4 ± 10.3 |
|  | TH | 19.7 (9.5) | 218.7 (22.3) | 124.9 (28.9) | 58.9 (36.5) | 34.1 (16.4) | 41.6(15.2) | 48.1 (12.5) |
|  | TH+EV | 21.7 (10.9) | 224.8 (17.9) | 136.3 (16.5) | 61.7 (18.7) | 39.7 (18.2) | 45.3 (20.5) | 45.1 (12.2) |

BE, base excess; TH, therapeutic hypothermia; TH+EV, TH plus edaravone; NT, normothermia. \* $p < 0.05$ , \*\* $p < 0.01$ , \*\*\* $p < 0.001$ vs. NT, <sup>##</sup> $p < 0.01$ vs.
TH+EV

**Fig 2.**
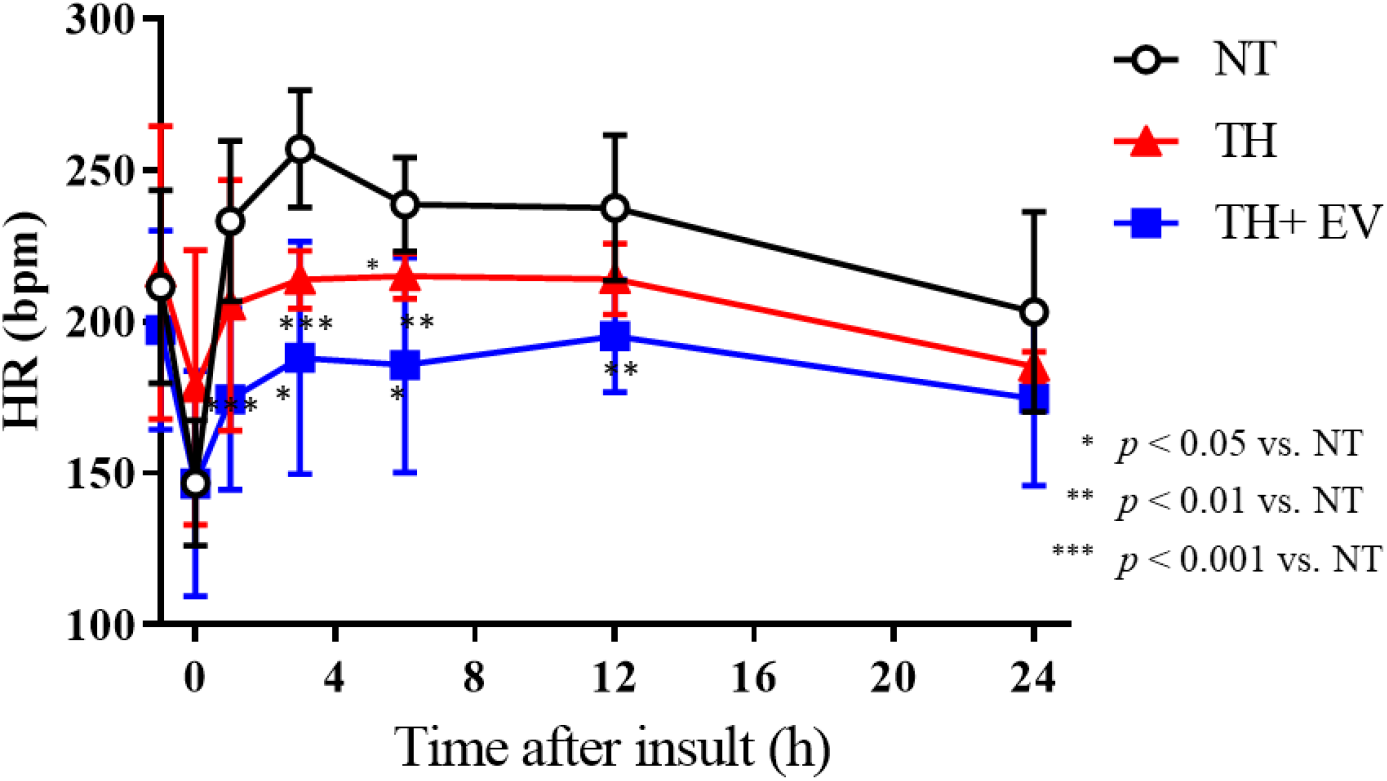
Time course of heart rate (HR) in three groups (NT, n=8; TH, n=7, TH+EV, n=6) at baseline, end of insult (0h), 3, 6, 12, 24 h after insult.

Before and during HI, rectal temperature was maintained in the normal range of 38.5–39.0 °C for piglets. In the hypothermic groups, the rectal temperature started to rapidly decrease 10 min after the resuscitation and reached 33.5 °C within 30 min. Thereafter, the temperature was well maintained (Fig 3).

**Fig 3.**
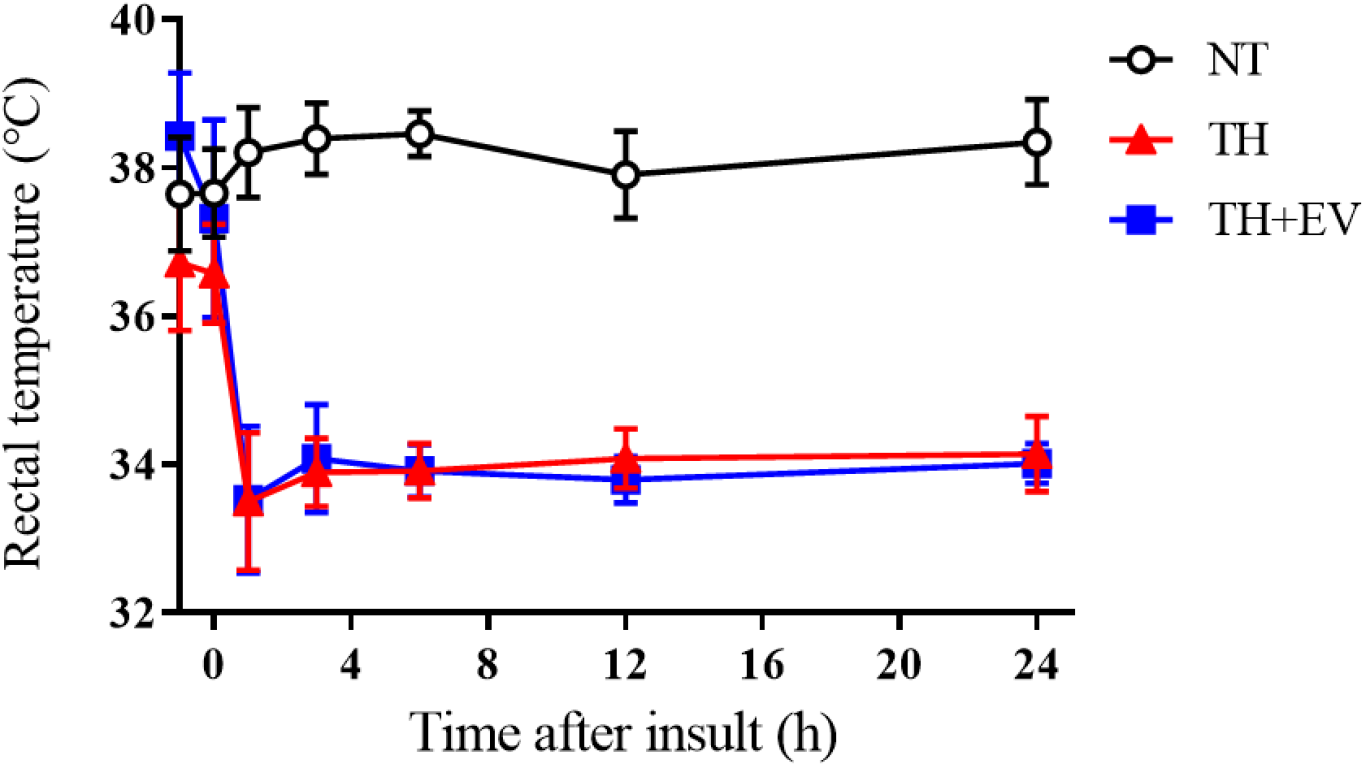
Time course of rectal temperature in three groups (NT, n=8; TH, n=7, TH+EV, n=6) at baseline, end of insult (0h), 3, 6, 12, 24 h after insult.

### Neurological assessment

At 5 days of recovery from HI, the mean (SD) neurological scores were 10.9 (5.7) in the HI-NT group, 17.9 (0.4) in the HI-TH group (*p* < 0.05 vs. HI-NT) and 15.0 (3.9) in the HI-TH+EV group (Fig 4). The percentages of piglets that could walk on day 5 after the insult were 25% (2 out of 8) in the NT group, 75% (6 out of 7) in the TH group and 40% (4 out of 6) in the TH+EV group.

**Fig 4.**
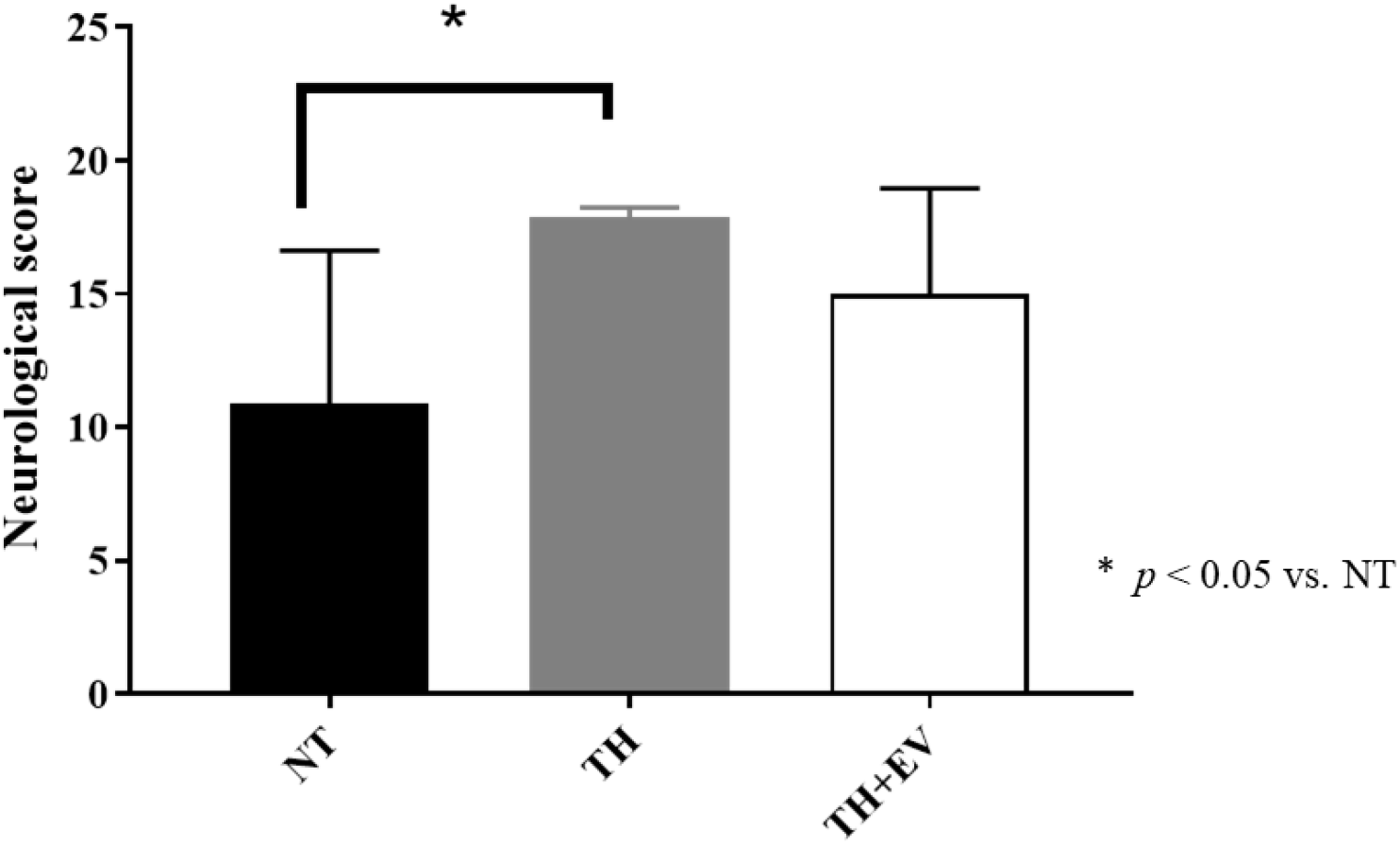
Neurological score at day5 after insult in three groups (HI-NT, n=8; HI-HT, n=8; HI-HT+EV, n=6)

### Histopathological results

Although the HI-TH group showed less damage in the cortex than the other groups, the HI-TH+EV group did not show less damage than the other groups (Fig 5a and b). There were no differences in damage among the three groups in the hippocampus and cerebrum (Fig 5c and d).

**Fig 5.**
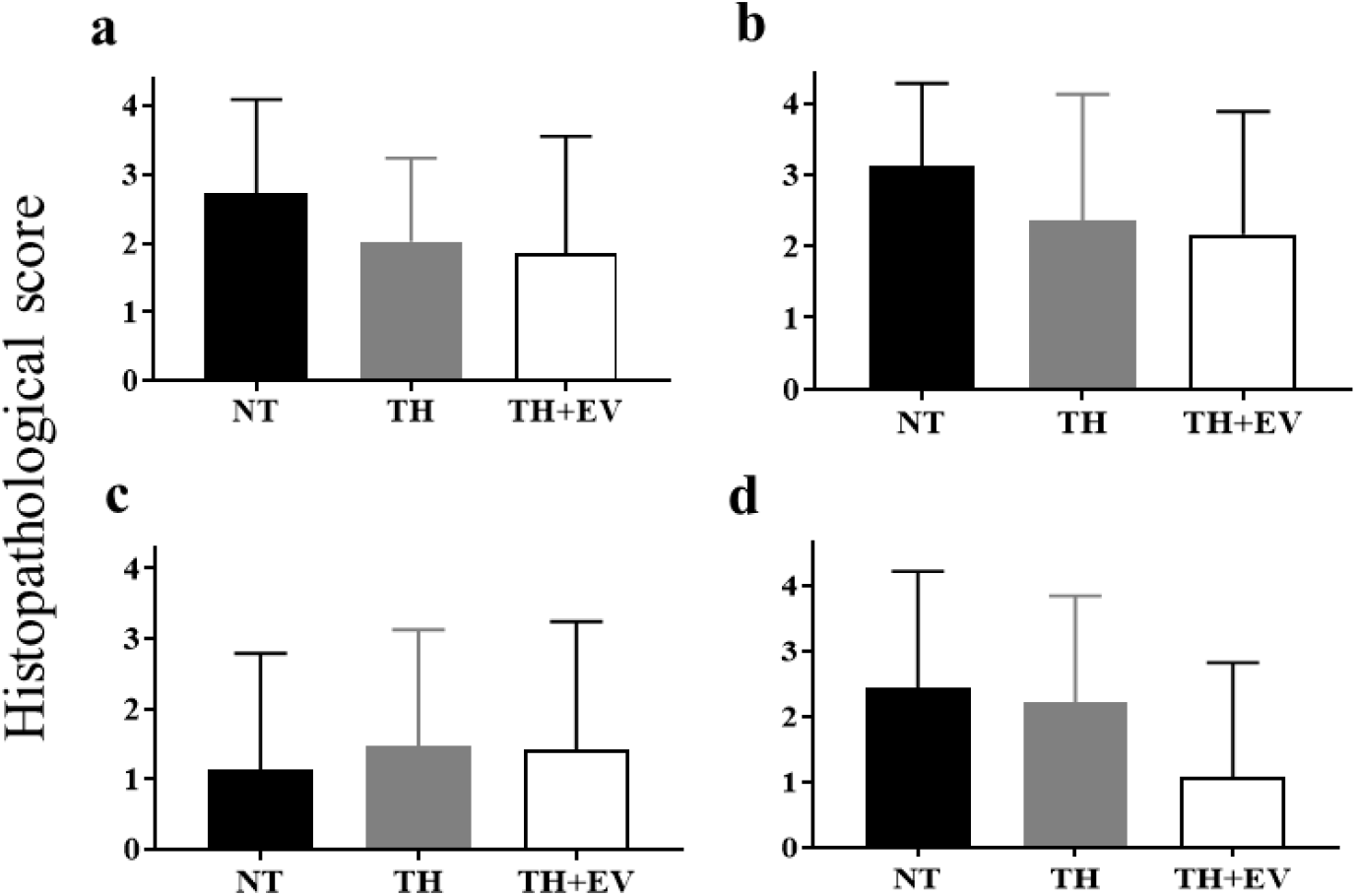
Histological scores among three groups; HI-NT, HI-HT and HI-HT+E in cortical GM(A), subcortical WM(B), CERE(C) and CA-1 of HIPP (D) (Means ± SD)

## Discussion

To our knowledge, this is the first report using a perinatal large-animal model to investigate the ability of intravenous administration of edaravone plus TH to protect against brain injury. The TH group showed an improved neurological score and walking ability on day 5 after the insult. Nevertheless, the confirmed the limited neuroprotective ability of edaravone plus TH. However, because many previous studies have reported the efficacy of edaravone for ameliorating brain injury in adult and neonatal animal models (Table 2), several possibilities need to be discussed to interpret this lack of efficacy.

**Table 2.** Summary of edaravone therapies in neonatal animal models after HI insult

|  | Animal | Route | Dosage and duration | Outcomes |
| --- | --- | --- | --- | --- |
| Ikeda et al., 2002<sup>16</sup> | 7-day-old rat | i.p. | 3, 6 and 9 mg/kg, 2 doses (second given 30 min after the first one) | Dose-dependent effectiveness (most effective with 9 mg/kg) |
| Yasuoka et al., 2004<sup>14</sup> | 7-day-old rat | i.p. | 3 mg/kg every 12 h until animal sacrifice (24 h, 48 h and 7 days) | Repeated injection reduced both necrosis and apoptosis |
| Noor et al., 2005a, b<sup>15</sup> | 7-day-old rat | i.p. | 9 mg/kg every 24 h until day 2, 5 or 10 | Two-day treatment with EV showed the best effect |
| Takizawa et al., 2009<sup>13</sup> | 7-day-old rat | i.p. | 9 mg/kg every 24 h until animal sacrifice | Brain damage reduced 48 h after insult |
| Ni et al., 2011<sup>17</sup> | 3–7-day-old piglet | i.v. | 3 mg/kg followed by 1.5 mg/kg/h for 5.5 h | Partial neuroprotection in striatum |

6 i.p., intraperitoneally; i.v., intravenously

The first possibility is that optimal efficacy is influenced by the dosage, number, interval and route (e.g., intravenous or intraperitoneal) of drug administration. The method for edaravone administration used in the present study was based on previous reports showing the efficacy of this method in the neonatal rat. In Yasuoka et al. [14], two doses of 3 mg/kg given to neonatal rat were found to be neuroprotective at only 24 h, but not 7 days, after the insult. Thus, we speculated that two doses were insufficient to maintain neuroprotection for long periods. Therefore, we increased the number of doses to six over 3 days. In a previous piglet study, Nii et al. [17] reported that intravenous edaravone 3 mg/kg plus 1.5 mg/kg/h for 5.5 h after an insult (total, 11.5 mg/kg) protected against brain injury. Positive results were obtained in 7-day-old rats given two daily intraperitoneal injections of 9 mg/kg [22] and 9-day-old Harlequin mice given two intraperitoneal injections of 10 mg/kg [23]. However, we administered 3 mg/kg every 12 h for 3 days after the HI insult. Thus, the dose in this study might not be enough to maintain the neuroprotective effects and our controversial results might be due to this dose difference.

In addition, referring to the administration of edaravone, a few reports have identified adverse effects in patients, such as acute renal failure [24] and fulminate hepatitis [25]. Given these adverse effects, the use of edaravone at extremely high doses might not be clinically useful. In normal adult humans, edaravone is well tolerated following single or multiple doses. Edaravone shows a short half-life in the range of 0.15– 5.16 h depending on the administration regimen [26]; otherwise, there is no evidence for the precise half-life of edaravone in the neonate. Future work is required to determine the half-life of edaravone to reveal the optimal administration methods for the newborn piglet.

The second possibility is that edaravone may not be effective under TH conditions. We expected that edaravone plus TH would show more additive neuroprotective effects than TH alone. However, in this study, TH alone clearly improved neurological outcomes compared with edaravone plus TH. This finding is compatible with those of many previous studies. Many previous animal and clinical reports suggested that TH had neuroprotective effects in adult stroke and neonatal HIE [27–29]. Edaravone has been shown to have neuroprotective properties in animal studies such as neonatal rats and young piglets under normothermia. We speculate that TH may reduce the efficacy of edaravone for brain injury. However, the mechanism of this effect is unclear. Shibuta et al. [30] reported that the temperature variations alter edaravone-induced neuroprotection against hypoxia-induced neurotoxicity and that temperature potentially determines whether edaravone confers a neuroprotective effect.

Another consideration is that TH is thought to protect the brain via multiple mechanisms, and it may thus be difficult to demonstrate the statistical significance of additional protection by another therapy. Because TH showed clear efficacy for neurological outcomes and improvement in walking in this study, it may have been complicated to prove the additional benefit of edaravone.

A limitation of this work is that we could not explore the mechanism explaining why edaravone could not improve neurological outcomes using our limited data analysis. Recent mechanistic research has suggested that edaravone suppresses delayed neuronal death [31], counteract microglia-induced neurotoxicity [32] and reduce long-term inflammation, not only via antioxidant activity [33]. In future work, we will reveal the mechanisms involved by examining cytokine profiles or other histological immunostains.

## Conclusion

We conclude that intravenous administration of edaravone plus TH for brain injury has no improvement on neurological outcome in the newborn piglet. Other methods of drug administration are necessary to address the efficacy of edaravone plus TH for brain injury in newborns.

## Acknowledgements

We thank the medical students of the Faculty of Medicine Kagawa University, Kagawa, Japan, who helped with this study.

## Supporting information

**S1 Fig Changes over time in the mean arterial blood pressure (MABP) in the three groups (NT, n = 8; TH, n = 7; TH+EV, n = 6) at baseline, end of insult (0 h) and 3, 6, 12 and 24 h after the insult.**

MABP was higher within 3 h after the insult in the TH (1 h, *p* < 0.05 vs NT) and TH+EV groups compared with the NT group. Values are shown as the mean ± SD. ^*^*p* < 0.05, ^**^*p* < 0.01, ^***^*p* < 0.001 vs. NT.

**S2 Fig Changes over time in the heart rate (HR) (b) and rectal temperature (c) in the three groups (NT, n = 8; TH, n = 7; TH+EV, n = 6) at baseline, end of insult (0 h) and 3, 6, 12 and 24 h after the insult.**

HR was significantly lower in the TH (1, 3, 6, 12 h) and TH+EV (3, 6 h) groups than in the NT group after the insult. Values are shown as the mean ± SD. ^*^*p* < 0.05, ^**^*p* < 0.01, ^***^*p* < 0.001 vs. NT.

**S3 Fig Changes over time in the rectal temperature (c) in the three groups (NT, n = 8; TH, n = 7; TH+EV, n = 6) at baseline, end of insult (0 h) and 3, 6, 12 and 24 h after the insult.**

**S4 Fig Neurological scores on day 5 after the insult in the three groups (NT, n = 8; TH, n = 7; TH+EV, n = 6).** The neurological score on day 5 was significantly higher in the TH group than in the NT group (*p* < 0.05).

**S5 Fig Histological scores on day 5 after the insult among the NT, TH and TH+EV groups in cortical grey matter (GM) (a), subcortical white matter (WM) (b), CA1 of the hippocampus (HIPP) (c) and the cerebrum (CERE) (d) (mean ± SD).**

**S1 Table. Mean (SD) values of arterial pHa, PaCO_2_, PaO_2_, base excess, blood glucose and lactate before, at the end of (0 h) and 1, 3, 6, 12 and 24 h after hypoxic-ischaemic insult in the three groups**

**S2 Table. Summary of edaravone therapies in neonatal animal models after HI insult**

## References

1. Liu L, Oza S, Hogan D, Perin J, Rudan I, Lawn JE, et al. Global, regional, and national causes of child mortality in 2000-13, with projections to inform post-2015 priorities: an updated systematic analysis. Lancet. 2015; 385: 430–440.

2. Tsuda K, Mukai T, Iwata S, Shibasaki J, Tokuhisa T, Ioroi T, et al. Therapeutic hypothermia for neonatal encephalopathy: a report from the first 3 years of the Baby Cooling Registry of Japan. Sci Rep. 2017; 7: 39508.

3. Gluckman PD, Wyatt JS, Azzopardi D, Ballard R, Edwards AD, Ferriero DM, et al. Selective head cooling with mild systemic hypothermia after neonatal encephalopathy: multicentre randomised trial. Lancet. 2005; 365: 663–670.

4. Jacobs SE, Berg M, Hunt R, Tarnow-Mordi WO, Inder TE, Davis PG. Cooling for newborns with hypoxic ischaemic encephalopathy. Cochrane Database Syst Rev. 2013; Cd003311.

5. Watanabe T, Yuki S, Saito K, Sato S. Jpn Pharmacol Ther. 1997; 25: 181–187.

6. Watanabe T, Yuki S, Egawa M, Nishi H. Protective effects of MCI-186 on cerebral ischemia: possible involvement of free radical scavenging and antioxidant actions. J Pharmacol Exp Ther. 1994; 268: 1597–1604.

7. Ochi H, Morita I, Murota S. Mechanism for endothelial cell injury induced by 15-hydroperoxyeicosatetraenoic acid, an arachidonate lipoxygenase product. Biochim Biophys Acta. 1992; 1136: 247–252.

8. Yamamoto T, Yuki S, Watanabe T, Mitsuka M, Saito KI, Kogure K. Delayed neuronal death prevented by inhibition of increased hydroxyl radical formation in a transient cerebral ischemia. Brain Res. 1997; 762: 240–242.

9. Shichinohe H, Kuroda S, Yasuda H, Ishikawa T, Iwai M, Horiuchi M, et al. Neuroprotective effects of the free radical scavenger Edaravone (MCI-186) in mice permanent focal brain ischemia. Brain Res. 2004; 1029: 200–206.

10. Lapchak PA. A critical assessment of edaravone acute ischemic stroke efficacy trials: is edaravone an effective neuroprotective therapy? Expert Opin Pharmacother. 2010; 11: 1753–1763.

11. Ferriero DM. Oxidant mechanisms in neonatal hypoxia-ischemia. Dev Neurosci. 2001; 23: 198–202.

12. Ferriero DM. Neonatal brain injury. N Engl J Med. 2004; 351: 1985–1995.

13. Takizawa Y, Miyazawa T, Nonoyama S, Goto Y, Itoh M. Edaravone inhibits DNA peroxidation and neuronal cell death in neonatal hypoxic-ischemic encephalopathy model rat. Pediatr Res. 2009; 65: 636–641.

14. Yasuoka N, Nakajima W, Ishida A, Takada G. Neuroprotection of edaravone on hypoxic-ischemic brain injury in neonatal rats. Brain Res Dev Brain Res. 2004; 151: 129–139.

15. Noor JI, Ueda Y, Ikeda T, Ikenoue T. Edaravone inhibits lipid peroxidation in neonatal hypoxic-ischemic rats: an in vivo microdialysis study. Neurosci Lett. 2007; 414: 5–9.

16. Ikeda T, Xia YX, Kaneko M, Sameshima H, Ikenoue T. Effect of the free radical scavenger, 3-methyl-1-phenyl-2-pyrazolin-5-one (MCI-186), on hypoxia-ischemia-induced brain injury in neonatal rats. Neurosci Lett. 2002; 329: 33–36.

17. Ni X, Yang ZJ, Carter EL, Martin LJ, Koehler RC. Striatal neuroprotection from neonatal hypoxia-ischemia in piglets by antioxidant treatment with EUK-134 or edaravone. Dev Neurosci. 2011; 33: 299–311.

18. Nakamura S, Kusaka T, Yasuda S, Ueno M, Miki T, Koyano K, et al. Cerebral blood volume combined with amplitude-integrated EEG can be a suitable guide to control hypoxic/ischemic insult in a piglet model. Brain Dev. 2013; 35: 614–625.

19. Nakamura S, Kusaka T, Koyano K, Miki T, Ueno M, Jinnai W, et al. Relationship between early changes in cerebral blood volume and electrocortical activity after hypoxic-ischemic insult in newborn piglets. Brain Dev. 2014; 36: 563–571.

20. Jinnai W, Nakamura S, Koyano K, Yamato S, Wakabayashi T, Htun Y, et al. Relationship between prolonged neural suppression and cerebral hemodynamic dysfunction during hypothermia in asphyxiated piglets. Brain Dev. 2018; 40: 649–661.

21. Thoresen M, Haaland K, Løberg EM, Whitelaw A, Apricena F, Hankø E, et al. A piglet survival model of posthypoxic encephalopathy. Pediatr Res. 1996; 40: 738–748.

22. Noor JI, Ikeda T, Mishima K, Aoo N, Ohta S, Egashira N, et al. Short-term administration of a new free radical scavenger, edaravone, is more effective than its long-term administration for the treatment of neonatal hypoxic-ischemic encephalopathy. Stroke. 2005; 36: 2468–2474.

23. Zhu C, Wang X, Huang Z, Qiu L, Xu F, Vahsen N, et al. Apoptosis-inducing factor is a major contributor to neuronal loss induced by neonatal cerebral hypoxia-ischemia. Cell Death Differ. 2007; 14: 775–784.

24. Hishida A. Determinants for the prognosis of acute renal disorders that developed during or after treatment with edaravone. Clin Exp Nephrol. 2009; 13: 118–122.

25. Abe M, Kaizu K, Matsumoto K. A case report of acute renal failure and fulminant hepatitis associated with edaravone administration in a cerebral infarction patient. Ther Apher Dial. 2007; 11: 235–240.

26. Shibata H, Shigenori A, Izawa M, Murksaki M, Takamatsu Y, Izawa O, et al. Phase I clinical study of MCI-186 (Edaravone, 3-methyl-1-phenyl-2-pyrazolin-5-one) in healthy volunteers: safety and pharmacokinetics of single and multiple administrations. Jpn J Clin Pharmacol Ther. 1998; 29: 863–876.

27. Thoresen M, Penrice J, Lorek A, Cady EB, Wylezinska M, Kirkbride V, et al. Mild hypothermia after severe transient hypoxia-ischemia ameliorates delayed cerebral energy failure in the newborn piglet. Pediatr Res. 1995; 37: 667–670.

28. Azzopardi DV, Strohm B, Edwards AD, Dyet L, Halliday HL, Juszczak E, et al. Moderate hypothermia to treat perinatal asphyxial encephalopathy. N Engl J Med. 2009; 361: 1349–1358.

29. Natarajan G, Pappas A, Shankaran S. Outcomes in childhood following therapeutic hypothermia for neonatal hypoxic-ischemic encephalopathy (HIE). Semin Perinatol. 2016; 40: 549–555.

30. Shibuta S, Varathan S, Kamibayashi T, Mashimo T. Small temperature variations alter edaravone-induced neuroprotection of cortical cultures exposed to prolonged hypoxic episodes. Br J Anaesth. 2010; 104: 52–58.

31. Lee BJ, Egi Y, van Leyen K, Lo EH, Arai K. Edaravone, a free radical scavenger, protects components of the neurovascular unit against oxidative stress in vitro. Brain Res. 2010; 1307: 22–27.

32. Banno M, Mizuno T, Kato H, Zhang G, Kawanokuchi J, Wang J, et al. The radical scavenger edaravone prevents oxidative neurotoxicity induced by peroxynitrite and activated microglia. Neuropharmacology. 2005; 48: 283–290.

33. Zhang W, Sato K, Hayashi T, Omori N, Nagano I, Kato S, et al. Extension of ischemic therapeutic time window by a free radical scavenger, Edaravone, reperfused with tPA in rat brain. Neurol Res. 2004; 26: 342–348.

